# *GPAT2* plays a role in the root cap cuticle formation of Arabidopsis that is not redundant to *GPAT4* and *GPAT8*

**DOI:** 10.1101/2025.04.25.650743

**Authors:** Alice Berhin, Aurore Guerault, Damien De Bellis, Christiane Nawrath

## Abstract

During the first days after germination, the root cap protects the seeding with its cuticle, which persists until the embryonic outer root cap cell layer is shed. The polyester constituting the delicate Arabidopsis root cap cuticle of the primary root has an atypical composition, containing unsubstituted very-long-chain fatty acids in addition to the oxygenated fatty acids typically found in cutin. The sn2-glycerol-3-phosphate acyltransferases GPAT4 and GPAT8, which possess an active phosphatase domain, synthesize monoacylglycerols and are crucial for incorporating oxygenated fatty acids into the root cap cutin. Here, we show that the incorporation of very-long-chain fatty acids into cutin requires the glycerol-3-phosphate acyltransferase GPAT2, which lacks an active phosphatase domain and produces lysophosphatidic acids. The GPAT2 knockout alters the properties of the root cap leading to increased mucilage deposition but lower barrier properties. Furthermore, we demonstrate that the long-chain CoA synthetase LACS2 is required solely for the incorporation of oxygenated fatty acids into cutin, similar to GPAT4 and GPAT8. Consequently, two largely non-redundant pathways contribute to root cap cuticle formation in the primary root: both the LACS2/GPAT4/GPAT8- and the GPAT2-dependent pathways contribute non-redundantly to its barrier and surface properties. LACS2 also plays an important role in the formation of the root cap cuticle in emerging lateral roots, whereas GPAT2 has a minor function in this process. These findings underscore the diverse mechanisms by which cuticle synthesis is fine-tuned across plant organs and developmental stages to achieve specialized barrier and surface properties.

## Introduction

Plants have adapted to variable environmental conditions on land by developing specialized cell wall modifications with barrier properties on the surfaces of their organs. A hydrophobic cuticle forms during embryogenesis, covering the embryo and playing a crucial role in plant development by preventing organ adhesion and fusion. Upon germination the cuticle of the shoot strengthens and expands with the growth of the plant continuously covering the epidermal tissues (Ingram and Nawrath 2017; De Giorgi et al. 2021). The cuticle covering the root cap however is shed along with the embryonic root cap cell layer when it is renewed (Berhin et al. 2019).

This root cap cuticle (RCC) is a delicate structure present on the outer layer of columella cells and the lateral root cap cells. It functions as a diffusion barrier protecting against unfavorable mineral and osmotic conditions (Berhin et al. 2019), and influences mucilage deposition on the root cap surface (Liu et al. 2024). Furthermore, the RCC covers also emerging lateral roots, acting as diffusion barrier and promoting their outgrowth from the primary root (Berhin et al. 2019).

The RCC has an electron-opaque ultrastructure resembling the ultrastructure of the Arabidopsis leaf cuticle (Nawrath et al. 2013; Berhin et al. 2019). The primary structural component of the Arabidopsis leaf cuticle is a cutin, which is rich in unsaturated dicarboxylic acids (DCA), particularly C18:2 DCA (Li-Beisson et al. 2013). While C18:2 DCA is also the main constituent of the RCC cutin, it contains in addition considerable amounts of saturated and monounsaturated very-long-chain fatty acids (VLCFAs) with 26 and 28 carbons, which is unusual for cutin (Berhin et al. 2019).

The formation of cutin precursors involves several enzymatic steps, including the CoA-activation of fatty acids by long-chain acyl-CoA synthetases (LACSs), oxygenation by cytochrome P450 monooxygenases, and transfer to glycerol-3-phosphate by glycerol-3-phosphate acyltransferases (GPATs). Additional modifications may be carried out by BAHD acyltransferases, such as DEFECTIVE IN CUTICULAR RIDGES (DCR). After being exported to the apoplast via ABCG transporters, cutin is polymerized by cutin synthases of the GELP-family, with additional contributions from α/β hydrolases such as BODYGUARD (BDG) (Fich, Segerson, and Rose 2016).

The biosynthetic pathway of the RCC cutin has only been partially characterized. The reduction of both oxygenated and very-long-chain unsubstituted fatty acids in the ester-bond lipids of the root cap of the *dcr* and *bdg* mutants suggests that both types of fatty acids are genuine constituents of RCC cutin (Berhin et al. 2019). A critical step in cutin precursor formation is the acyl transfer of CoA-activated fatty acids to glycerol-3-phosphate, catalyzed by land-plant specific glycerol-3-phosphate acyltransferases (GPATs) (Yang et al. 2012). In Arabidopsis, GPAT4 and GPAT8 are essential for forming leaf and stem cutin rich in dicarboxylic acids (Li et al. 2007). *gpat4 gpat8* double mutants exhibit also reduced levels of dicarboxylic acids and oxygenated fatty acids in the RCC cutin of the primary root (Berhin et al. 2019) but the VLCFA content remains unchanged, raising questions about which GPATs are resonponsible for VLCFA integration into the RCC cutin.

The eight members of the land-plant-specific GPAT family exhibit *sn2*-regiospecificity for acyl transfer to glycerol-3-phosphate (Yang et al. 2012). GPAT4, GPAT6 and GPAT8 also possess phosphatase activity, enabling them to synthesize monoacylglycerols (MAGs) (Li et al. 2007; Li-Beisson et al. 2009). In contrast, the other GPAT family members have a mutated phosphatase domain and instead synthesize lysophosphatidic acids (LPAs). GPAT5 and GPAT7, closely related to GPAT4, GPAT6, and GPAT8, act primarily in suberin biosynthesis, a polymer related to cutin (Beisson et al. 2007; Gully et al. 2024). More distantly related, GPAT1, GPAT2 and GPAT3 play diverse roles. For example, GPAT1 is involved in tapetum formation and male fertility alongside GPAT6 (Zheng et al. 2003). Additionally, GPAT1 and GPAT2 function redundantly in auxin-signalling by synthesizing lysophosphatidic acids, which bind to PIN proteins for activation (Jia et al. 2022). However, no function of GPAT2 and GPAT3 in polyester synthesis has been identified so far (Yang et al. 2012).

Here, we demostrate that *GPAT2* is essential for RCC formation at the cap of the primary root. *GPAT2* knockout results in an altered RCC ultrastructure, increased permeability and significantly impacts mucilage deposition at the root cap of the primary root. Notably, the levels of VLCFAs and their monounsaturated derivatives are significantly reduced in RCC cutin. We show that *GPAT2* has non-redundant roles to *GPAT4* and *GPAT8* in RCC formation, which may be related to differences in substrate-specificty. Furthermore, we identify LONG-CHAIN ACYL-COA ACYLTRANSFERASE 2 (LACS2) as a key factor in RCC formation in both primary and outgrowing lateral roots. *LACS2* knockout selectively affects the incorporation of oxygenated fatty acids, similar to *GPAT4* and *GPAT8,* suggesting that *LACS2*, *GPAT4*, and *GPAT8* operate in a distinct pathway independent of *GPAT2*.

## Results

### *GPAT2* and *LACS2* are essential for RCC formation in primary roots

*GPAT4* and *GPAT8* are crucial for cutin synthesis of the RCC; however, ester-bound lipids are reduced by only 50% in the *gpat4 gpat8* mutant in the primary root, suggesting involvement of additional *GPATs* in RCC cutin synthesis. Expression analysis of the additional land-plant specific GPATs (*GPAT1*/*2*/*3*/*5*/*6*/*7*) in root cap cells was performed in 2-day-old seedlings using pGPATx::nlsGFP-GUS transgenic Arabidopsis lines.

*GPAT2* and *GPAT3* expression was detected in the outermost root cap cell layer and epidermal cells of the roots, while *GPAT1* exhibited only weak expression in outermost columella cells but was visible in lateral root cap and root meristem cells (Figure S1A). *GPAT5*, *GPAT6,* and *GPAT7* were not expressed in the root cap (Figure S1A).

To assess a possible role of GPAT1/2/3 in RCC formation, 2-day-old seedlings of *gpat1, gpat2*, and *gpat3* were stained with Fluorol Yellow (FY) to detect aliphatic polyester depositions in the root. In *gpat1* and *gpat3* (Figure S2A), FY staining was detected in the outher root cap cell layer indicating the presence of a RCC. In contrast, *gpat2* mutants showed reduced FY staining, limited to columella cells, in the primary root (Figure 1A and Figure S2A). This phenotype was rescued by expressing *GPAT2* in the *gpat2-1* mutant (Figure S2A), confirming the role *GPAT2* in RCC formation.

**Figure 1.**
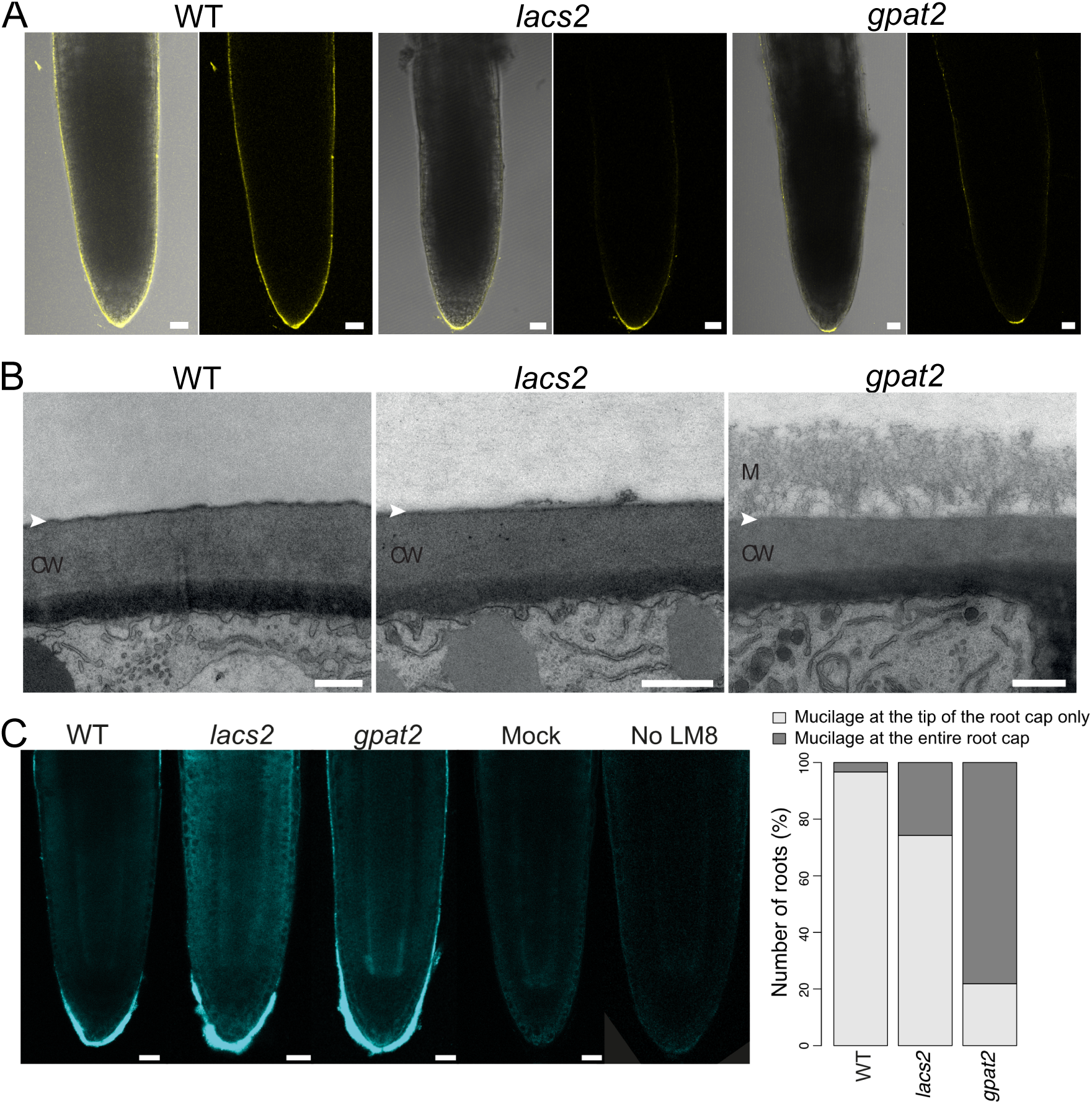
Deficiencies in root cap cuticle formation of *gpat2* and *lacs2* seedings at the primary root. (A) Reduced amounts of polyester in the RCC of primary roots as visualized by Fluorol yellow staining. Medium view of the root cap of 2-day-old wild type (WT) plants as well as *gpat2* and *lacs2* mutants after Fluorol Yellow staining (left side: overlays between bright field and fluorescence images; right side: fluorescence only). Scale bar = 20 µm. (B) Ultrastructure of the cuticle at the root cap of primary roots. Transmission electron micrographs showing the cell wall and cuticle of the outermost root cap cell layer of primary roots of 2-day-old WT plants as well as *lacs2* and *gpat2* mutants. Scale bar = 500 nm; CW, cell wall; M, mucilage; white arrowhead: expected position of the cuticle. (C) Evaluation of the mucilage deposition on the surface of root cap cells of the 2-day-old seedlings of diperent genotypes. Median view of a representative confocal fluorescence micrograph on the left and quantification of the mucilage deposition on the right. Mock, no antibodies. Scale bar = 20 µm.

*LACS2,* known for its role in cutin synthesis in leaves and other shoot organs (Bessire et al. 2007; Zhao et al. 2019), was also tested for its involvement in RCC formation. *LACS2* had been shown previously to be expressed in root cap cells (Lu et al. 2009)*. Lacs2* mutants exhibited a strongly reduced FY staining in two day-old seedlings (Figure 1A).

Thus, *LACS2* and *GPAT2* are essential for the formation of the RCC in the primary root of 2-day-old seedlings.

### *GPAT2* and *LACS2* knockouts affect the RCC ultrastructure of primary roots differently

Transmission electron microscopy of 2-day-old Arabidopsis roots revealed an electron-dense RCC in wild-type plants (Figure 1B) (Berhin et al. 2019). In *gpat2* mutant plants, the RCC was nearly absent, often accompagnied by an extensive presence of mucilage. In *lacs2* mutant plants, the RCC was thinner than in wild type (Figure 1B), but mucilage was rarely observed on it surface (Figure 1B).

Quantification of the presence of mucilage at the RCC in 2-day-old seedlings using the LM8 antibody recognizing xyloglucane revealed more frequent mucilage presence at lateral root cap cells in *gpat2* (78%) than in *lacs2* (26%) or wild-type (3%) (Figure 1C).

The results suggest that while both *GPAT2* and *LACS2* contribute to RCC formation, *GPAT2* plays a broader role for the surface properties of the root cap.

### Different contributions of *GPAT2* and *LACS2* to RCC cutin synthesis and barrier properties

Chemical analysis of polyester monomers in 2-day-old seedlings (having no endodermal suberin; Berhin et al., 2019) showed that the *GPAT2* knockout reduced C18:2 DCA by approximately 30% and decreased C26/C28 VLCFAs by 92%, leading to a 68% reduction in unsubstituted VLCFAs compared to wild type (Figure 2A).

**Figure 2.**
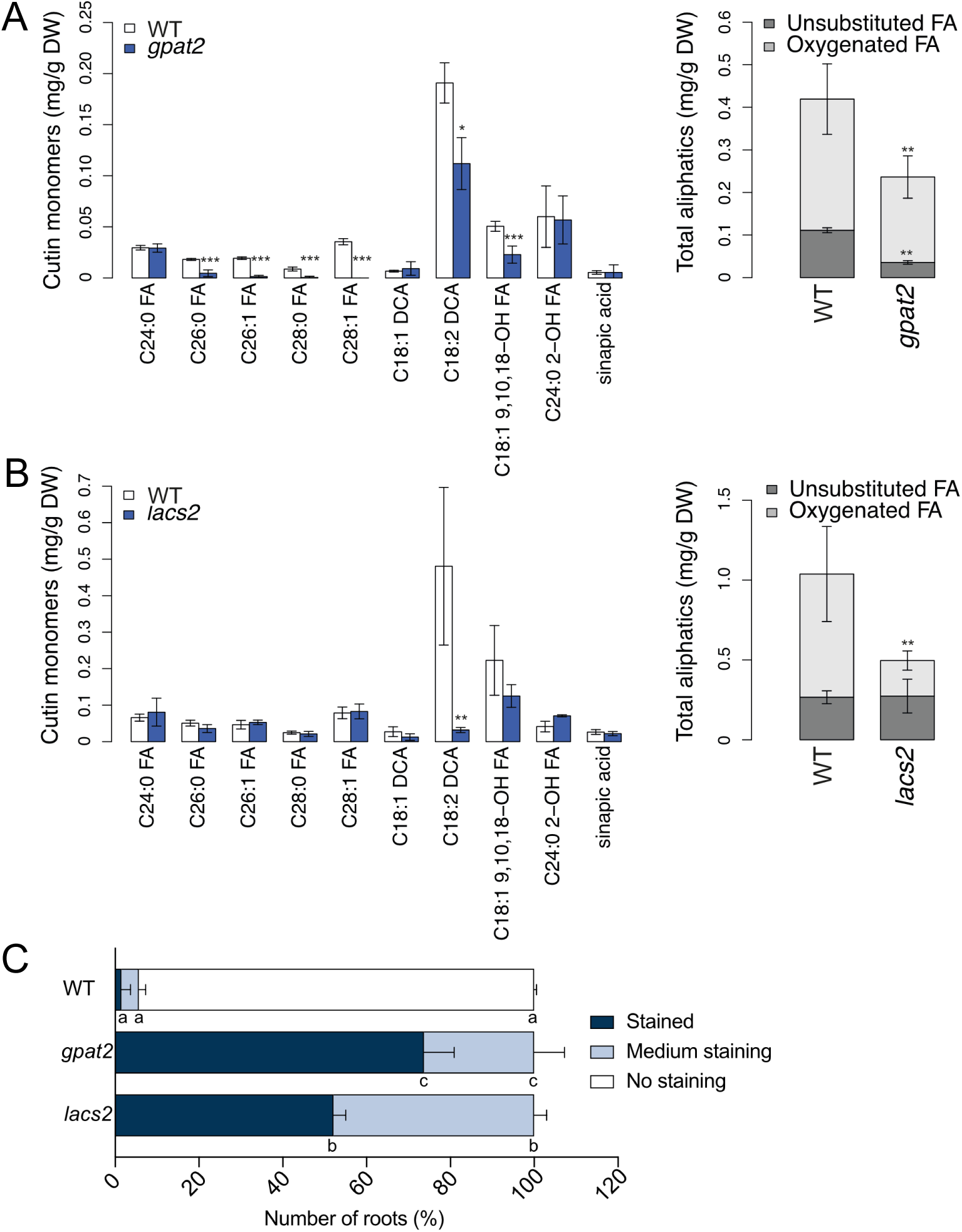
Reduced amounts of cutin in the root cap cuticle of the primary root and increased cuticle permeability of *gpat2* and *lacs2* seedlings. The composition of cutin in primary roots of 2-day-old seedlings of (A) *gpat2* and (B) *lacs2* mutants in comparison to the wild type (WT). Quantification of aliphatic and aromatic ester-bond cutin monomers obtained after transesterification. Left graph shows the principal cutin monomers and right graph shows the total of evaluated aliphatic compounds grouped by substance classes. Values represent the means ± SD, n = 3–4. Asterisks denote significant differences to as determined by Student’s t-test: ***p < 0.001; **p < 0.01. FA, fatty acid; DCA, dicarboxylic acid; DW, dry weight; (C) Assessment of the permeability of the root cap cuticle of the primary root by Toluidine Blue staining in *gpat2* and *lacs2* mutants in comparison to the wild type. 20 to 30 roots were tested per replicates. Values represent the means of two repetitions and significant differences to wild type were assessed by ANOVA. Letters represent statistically significant differences between groups (p < 0.05).

The *LACS2* knockout reduced C18:2 DCA by 93% and total oxygenated fatty acids by 70%, while unsubstituted fatty acids remained unchanged (Figure 2B).

Toluidine Blue (TB) staining was used to assess cuticle permeabilty of the RCC in 2-day-old roots (Berhin et al. 2019). High TB penetration demonstrated a higher RCC permeability in *gpat2* and *lacs2* mutants compared to wild type, with *gpat2* showing a more pronounced effect (Figure 2C, Figure S2B). These findings point to a particular role of VLCFAs in forming stronger difusion barriers.

### *LACS2,* but not *GPAT2,* strongly affects the RCC deposition in emerging lateral roots

The emerging lateral root also has a cuticle at its cap (Berhin et al. 2019). *LACS2* had already been shown to be expressed at the root cap of the lateral root (Lu et al. 2009). Analysis of outgrowing lateral roots in 8-day-old transgenic pGPAT2::nls-GFP-GUS expressing Arabidopsis plants did not indicate *GPAT2* expression in the lateral root primordium and young emerging lateral root (Figure S1B). Nevertheless, the RCC of emerging lateral roots was examined in both *lacs2* and *gpat2* mutants.

Absence of FY staining indicated the lack of a functional RCC from the lateral root in *lacs2*, both from columella and lateral root cap cells. A FY staining surrounded the entire root cap in the *gpat2* mutant, nevertheless appearing weaker (Figure 3 A). We included the *gpat2* mutant in subsequent analyses of the RCC at the outgrowing lateral root to obtain a more detailed view.

**Figure 3.**
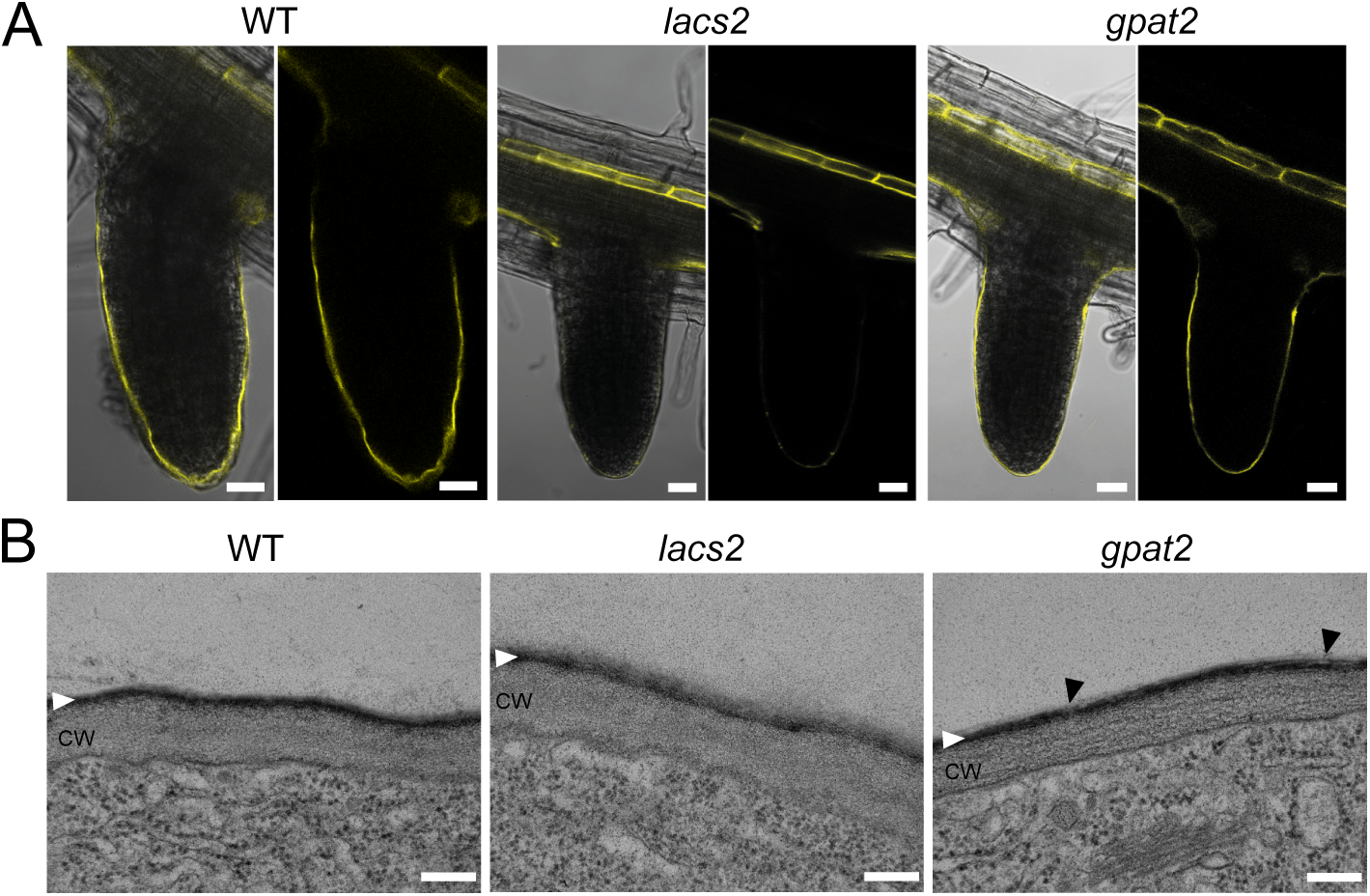
Stronger modifications of the root cap cuticle of outgrowing lateral roots in the *lacs2* than in the *gpat2* mutant. (A) Fluorol yellow staining of the root cap cuticle of outgrowing lateral roots in 8-day-old seedlings revealed strong impairment in polyester deposition in the *lacs2* mutant, but not in *gpat2* in comparison to the wild type (WT). Medium views of the root cap are shown (left side: overlay between bright field and fluorescence; right side: fluorescence only). Scale bar = 20 µm. (B) Ultrastructure of the cuticle of the root cap in outgrowing lateral roots. Transmission electron micrographs showing the cell wall and cuticle of the outermost root cap cell layer of lateral roots of 2-day-old WT plants as well as *lacs2* and *gpat2* mutants indicating ultrastructural modifications in the *lacs2* and *gpat2* mutant in comparison to the WT. Scale bar = 500 nm; CW, cell wall; white arrowhead, expected position of the cuticle, black arrowheads, structural impairments in the *gpat2* cuticle.

Transmission electron microscopy showed that the cuticle of the lateral root cap was more substantial than in primary roots in the wild type (Berhin et al. 2019). A more eroded and less contrasted ultrastructure of the cuticle in root cap cells was observed in the newly emerged lateral roots in *lacs2* in comparison to the wild type (Figure 3B). The cuticle in the *gpat2* mutant had similar dimensions and contrast to that of the wild type (Figure 3B). However, substructures were visible within the cuticle of *gpat2* that were not seen in the wild type indicating minor structural changes in the cuticle.

In summary, investigations of RCC deposition at outgrowing lateral roots of *lacs2* revealed a profound impact of the *LACS2* knockout on RCC formation. Possible contributions of *GPAT2* to the formation of RCC of the lateral root were less pronounced.

### Functional consequences of *LACS2* and *GPAT2* knockouts for the RCC in outgrowing lateral roots

To relate ultrastructure to properties, fluorescein diacetate (FDA), a dye that gets converted to a fluorescent compound after entering living cells, was used to assess RCC permeability of recently emerged lateral roots (Berhin et al. 2019; Barberon et al. 2016). The root cap cells of wild type showed no increased fluorescence in accordance with its intact cuticle (Figure 4A). FDA staining showed increased permeability in *lacs2,* but minimal changes in *gpat2* mutants (Figure 4A).

**Figure 4.**
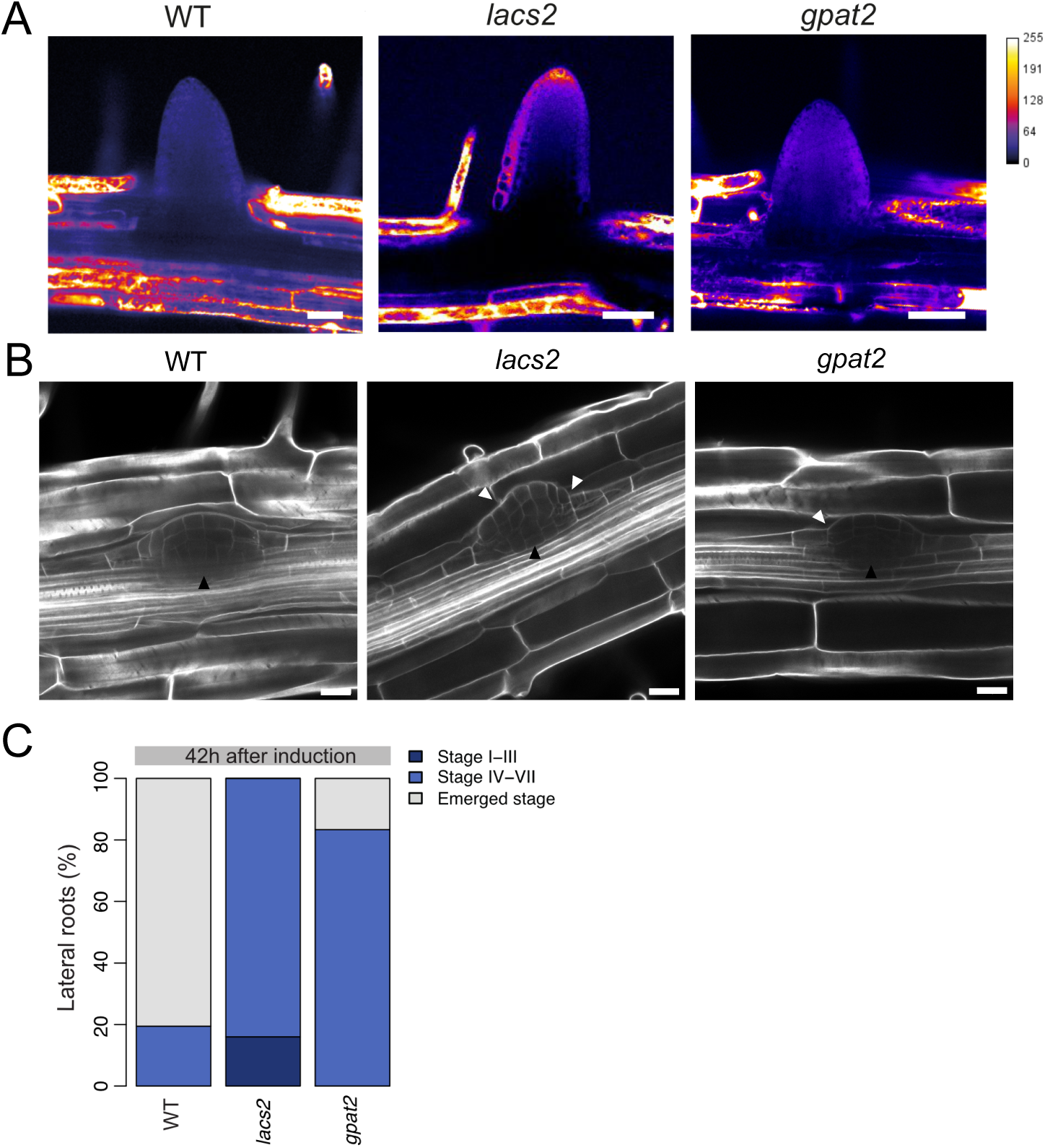
Stronger impact on properties of the outgrowing lateral root cuticle in the *lacs2* than in the *gpat2* mutant. (A) Penetration of the fluorescent cellular tracer fluorescein diacetate into root cap cells and meristematic cells of recently emerged lateral roots of 8-day-old seedlings of wild type (WT) plants and *lacs2* and *gpat2* mutants after 4 min of incubation. Relative intensity of the fluorescence is depicted by the color code. Scale bars = 50 μm. (B) Shape of the lateral root primordia in WT and *lacs2* having a strongly modified RCC at the emerging lateral root. Regular shape of a lateral root primordium of WT and deformed lateral root primordia having RCC modifications at the lateral root. Black arrowhead, lateral root primordium; white arrowhead, primordium deformation. Scale bars = 20 mm. (C) Stages of lateral root primordia 42 h after induction were evaluated in WT and in *lacs2* and *gpat2* at the lateral root primordium. Stages were determined as described in Casimiro et al. (2003): stage I–III, before breakage into the cortex; stage IV–VII, within the outer layers of the primary root; emerged, outside of the primary root.

Modified surface properties of the root cap cuticle at the lateral root may influence the outgrowth of the lateral root (Berhin et al. 2019). The outgrowing root formed a regular dome in the wild type, showing no indication of adhesion to surrounding tissues. In contrast, in the *lacs2* mutant, the RCC occasionally adhered strongly to other cells during lateral root outgrowth, leading to deformations of the emerging lateral root (Figure 4B). In the *gpat2* mutant, only slight adhesion effects were occasionally observed between the cuticle of the emerging lateral root and endodermal cells of the main root (Figure 4B).

None of the lateral roots of *lacs2* had emerged after 42 hours, whereas 80% of the wild-type roots had (Figure 4C). The delay in lateral root outgrowth in *gpat2* was less pronounced than in *lacs2*; only 20% of the lateral roots had emerged in *gpat2*, none were at the early lateral root developmental stage, as observed in the *lacs2* mutant (Figure 4C). The impact of cuticle impairments in the *lacs2* mutants on the lateral root emergence was also reflected in the shape of the outgrowing lateral root.

In summary, *LACS2* is critical for RCC integrity and lateral root emergence, while *GPAT2* plays a lesser role but still contributes to RCC properties of the outgrowing lateral root.

## Discussion

With the presented studies we extended our knowledge on root cap cuticle formation by revealing different roles of *GPAT2* and *LACS2* for the formation of the cuticle at the root cap of primary and recently-outgrowing lateral roots.

*GPAT2* plays a crucial role in root cap cuticle formation, in the young primary root and to a lower extent at the outgrowing lateral root. *gpat2* mutant RCC structure was strongly affected at the primary root but not at the lateral roots (Figure 1A-B, Figure 3A-B, Figure S1A-B, Figure S2A) leading to proportional permeability increases (Figure 2C, Figure 4A, Figure S2B) and lateral root emergence defects (Figure 4B-4C). Unlike other cuticles, the root cap cuticle of the primary root contains a high proportion of ester-bond VLCFAs, in addition to oxygenated fatty acids. The knockout of *GPAT2* drastically reduced the unusual C26/C28 VLCFAs, while the reduction of the typical oxygenated fatty acids of the cutin of the RCC in the primary root was moderate (Figure 2A). Esterified VLCFAs cannot contribute to the ramification of the lipid polyester network but act as polyester chain terminators. Their low polarity likely enhances water-repelling properties and may thus contribute to better sealing of the delicate cuticle, similar to wax compounds (Lewandowska, Keyl, and Feussner 2020). This may explain that the cuticle permeability in *gpat2* was higher than in *lacs2* (Figure 2C). An interesting difference in mucilage deposition was noted between the mutant genotypes, with the knockout of *GPAT2* facilitating deposition more strongly than the one of *LACS2* (Figure 1C), which is likely related to their respective RCC impairments (Liu et al. 2024).

The differences in how *GPAT2* and the redundantly acting *GPAT4* and *GPAT8* affect the ester-bound lipid composition and/or the structure may account for their non-redundant roles in RCC formation. GPAT2 and GPAT4/GPAT8 likely exhibit different substrate specificities impacting the ester-bond lipid composition differently. Furthermore, GPAT2 carries mutations in the intrinsic phosphatase domain synthetizing lysophosphatidic acids instead of monoacylglycerols (Yang et al. 2012). Recent studies have revealed ultrastructural differences in the suberin polymer, which are closely associated with the activity of the phosphatase domain, i.e. associated with the generation of lysophosphatidic acids by GPAT5 and GPAT7 (Gully et al. 2024). Therefore, it is possible that different cutin precursor types that are generated by GPAT2 and GPAT4/GPAT8 may also affect cutin structure in the RCC (Figure 5). Similarly, GPATs synthetizing different precursor types contribute also to sporopollenin formation, i.e. GPAT1 synthetizing lysophosphatidic acids and GPAT6 monoacylglycerols (Li et al. 2012).

**Figure 5.**
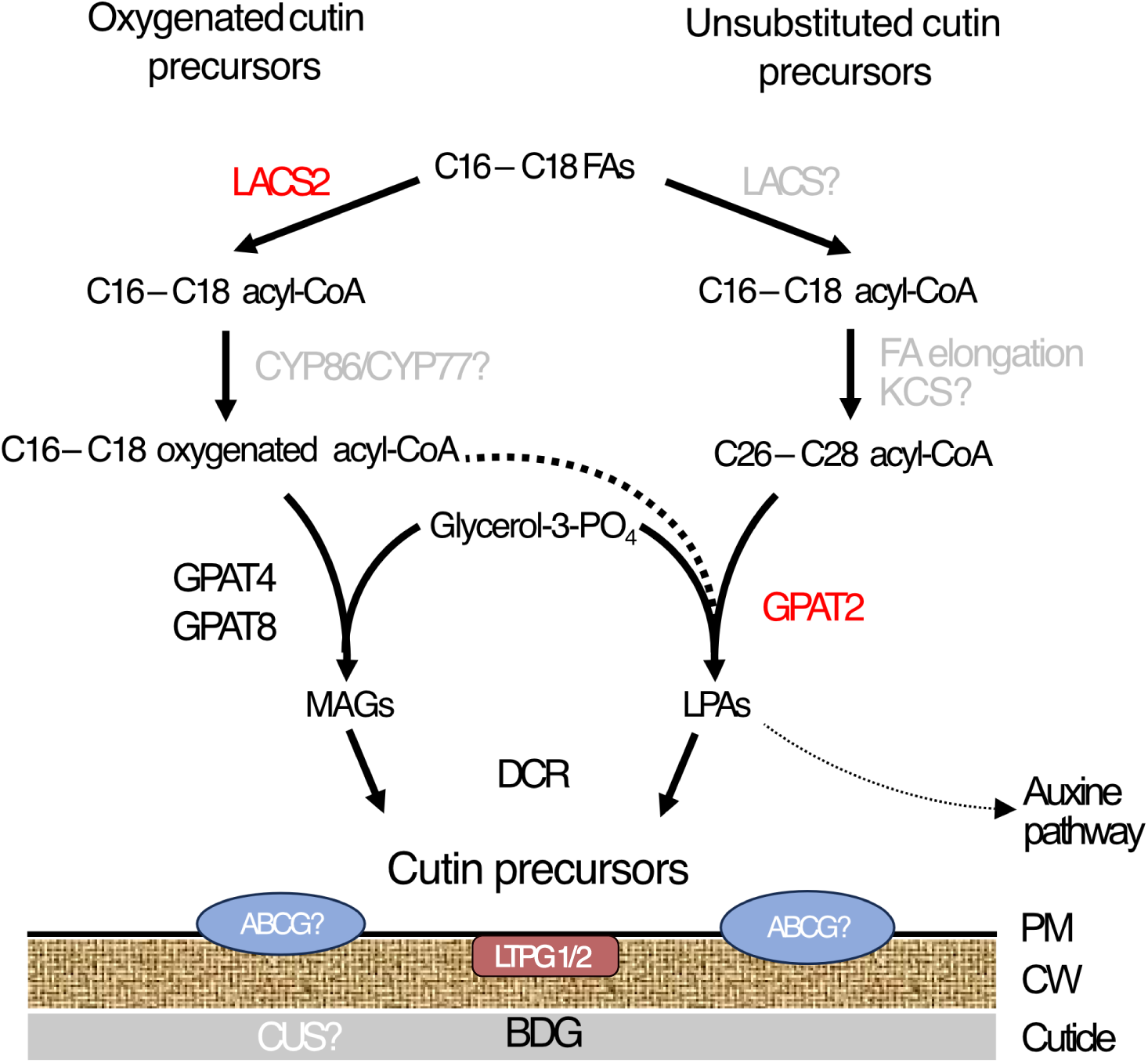
Schematic diagram summarizing the biosynthetic pathway of cutin of the root cap cuticle of primary roots. LACS2 and GPAT4/8 act in the branch of formation of oxygenated cutin precursors only, while GPAT2 has specific functions for the incorporation of very-long-chain fatty acids into cutin. Genes described in this study are indicated in red, genes characterized in root cap cuticle formation are indicated in black (Berhin et al. 2019), genes postulated are indicated in gray. Unknown isoforms are indicated by a question mark. Minor contributions to cutin biosynthesis at the RCC are indicated by a dashed line. Hypothesized contribution of LTPG1/2 to cutin formation at the root cap (Uemura et al. 2024). Potential contributions to other pathways are indicated by a thin dashed line (Jia et al. 2022).

Most enzymes involved in cutin and wax formation, including LACS2 and GPAT8, were localized to endoplasmic reticulum (ER) (Gidda et al. 2009; Weng et al. 2010). Although GPAT2 lacks a clear localization sequence, recent studies suggest mitochondrial localization (Jia et al. 2022), contrasting earlier findings (Zheng et al. 2003). Further research under physiological conditions is required to determine the localization of GPAT2, as well as GPAT1 and GPAT3. In mammals *sn1*-GPATs localize to both, the outer mitochondrial membrane and the ER membrane with the catalytic site in the cytosol (Huang et al. 2022). This raises the possibility for mitochondrial and ER-localization of GPATs with *sn2*-regiospecificity in plants, enhancing the number of regulatory mechanisms. GPAT1/GPAT2/GPAT3 might have a broader range of functions in plant physiology than GPAT4/GPAT8 since LPAs derived from GPAT1 and GPAT2 are necessary for PIN protein regulation and polar auxin transport (Jia et al. 2022), in addition to their contributions to the formation of polymers of the apoplast (this study, Zheng et al. 2003). In contrast, GPAT4/GPAT8 are required only for the formation of different barriers of the apoplast (Nawrath et al. 2013). Whether this is related to different localization remains to be discovered.

The *LACS2* knockout only reduces the amount of oxygenated fatty acids in cutin of various organs indicating a substrate specificity for oxygenated fatty acids (Figure 2B) (Bessire et al. 2007; Zhao et al. 2019). The resemblance in composition of the ester-bound lipid composition and the total cutin amount in the RCC between the *lacs2* mutant and the *gpat4 gpat8* double knockout (Berhin et al. 2019) suggest these enzymes function within the same cutin biosynthesis pathway branch (Figure 5). This resemblance might also imply metabolic channeling of acyl precursors for cutin synthesis, which is an important regulatory mechanism in plant metabolism that also occurs at the ER (Figure 5) (Kriechbaumer and Botchway 2018; Zhang and Fernie 2021). Further investigations are necessary to gain additional insights into the possible existence of a metabolon for cutin synthesis at the ER in plants.

Our characterization of the RCC in the outgrowing lateral root shows that *GPAT2* plays a more prominent role in the RCC formation in primary root than in lateral roots, while LACS2 is essential in both. The characterization of *gpat2* in outgrowing lateral roots exemplifies the current limitations in revealing differences in cuticle formation by histochemical staining and transmission electron microscopy. Assays to assess cuticle permeability are more sensitive in the detection of cuticle impairments, similar as more derived impact of cuticle alterations on the plant development, a phenomenon that had also been observed previously (Bessire et al. 2011; Fabre et al. 2016).

Our results show that the composition, structure and properties of the cuticle vary among different cell types of the root cap (columella and lateral root cap cells) and across different developmental stages (root caps of primary and outgrowing lateral roots). Similarly, the cuticles of aerial organs exhibit broad variations in their composition among organs and developmental stages (Nawrath et al. 2013; Fabre et al. 2016). Further studies will be necessary to better understand the relation between structure, composition and function in plant cuticles in general.

## Conclusions

Plants form cuticles on their organ surfaces that protect themselves from environmental stresses and provide appropriate surface properties for developmental processes. These cuticles exhibit diverse compositions and physical properties. Our discovery that *GPAT2*, encoding a GPAT that forms lysophosphatidic acid, contributes to cutin formation, alongside with *GPATs* responsible for the formation of monoacylglycerols, adds a new dimension to our understanding of how plants generate the wide variety of cuticular structures crucial for their survival. Additionally, our findings underscore the essential function of LACS2 in incorporating oxygenated precursors into cutin across different organs, potentially functioning specifically within the GPAT4/GPAT8-dependent biosynthetic pathway.

## Acknowledgements

We would like to thank Nasim Farahani Zayas for her contribution of the pGPAT1::nls GFP-GUS lines. The NASC stock center is thanked for providing seed stocks and vectors. Furthermore, we thank Christel Genoud and Arnaud Paradis for leading the Electron Microscopy Facility and the Imaging Facility of UNIL, respectively. This work was supported by the Swiss National Science Foundation (grant 31003A_170127 and 310030_188672 to CN).

## Author contributions

A.B. and A.G. performed experiments and data evaluation; D.D.B. performed electron microscopy studies; C.N. conceived the project, supervised the research, and wrote the article with contributions from all authors.

## Figures

**Figure S1.**
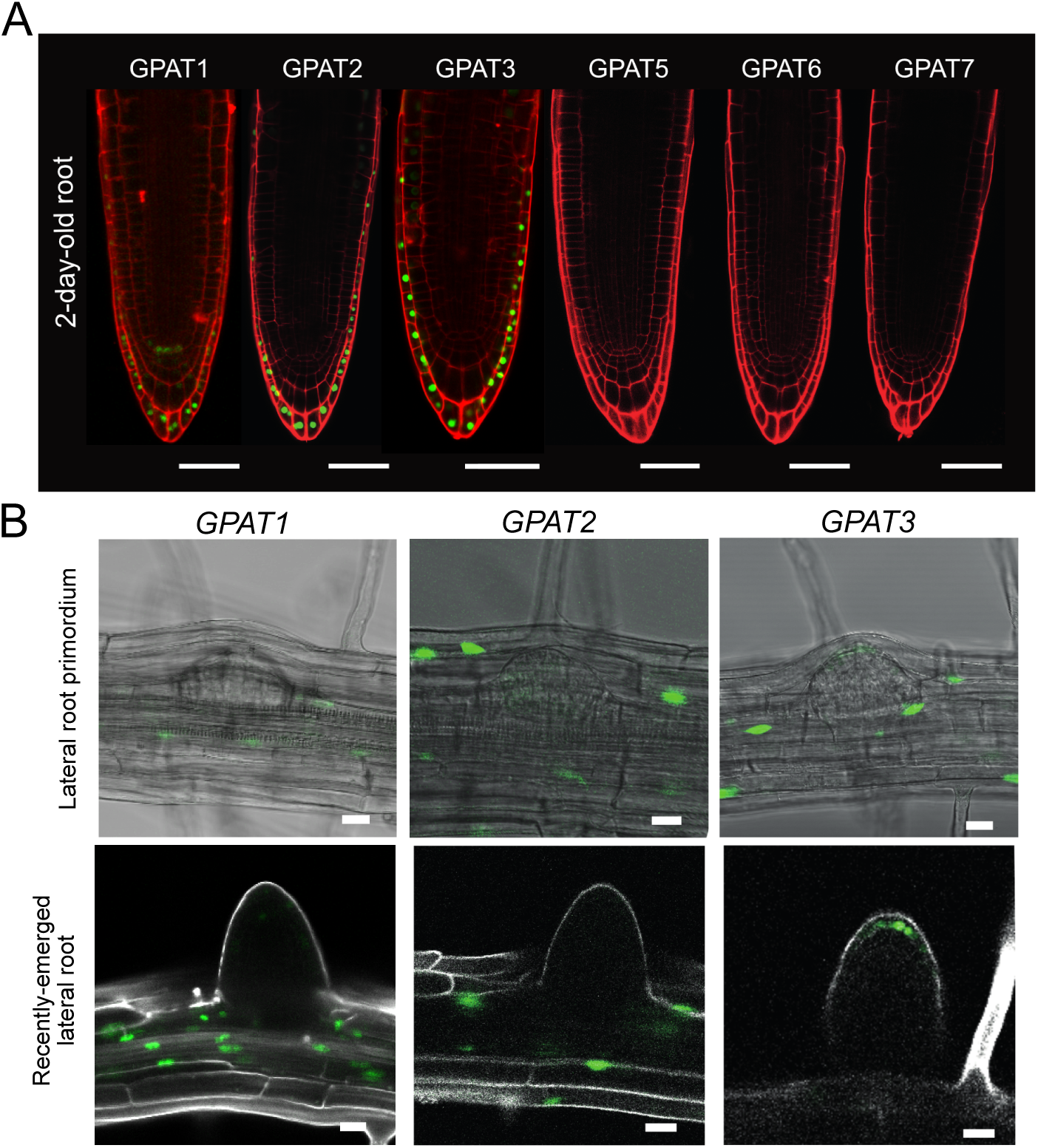
Expression of *GPATs* at the tip of the primary and outgrowing lateral roots. Confocal images of transgenic plants expressing pGPATx::nlsGFP-GUS in root cap cells of the primary root of 2-day-old seedlings (A) or in root cap cells of lateral roots of different stages of 8-day old roots (B). GFP fluorescence is shown in green. Cell outlines are visualized by propidium iodide staining in red (A), and Calcofluor white staining in white (B). Scale bars = 50 µm (A) and 20 µm (B).

**Figure S2.**
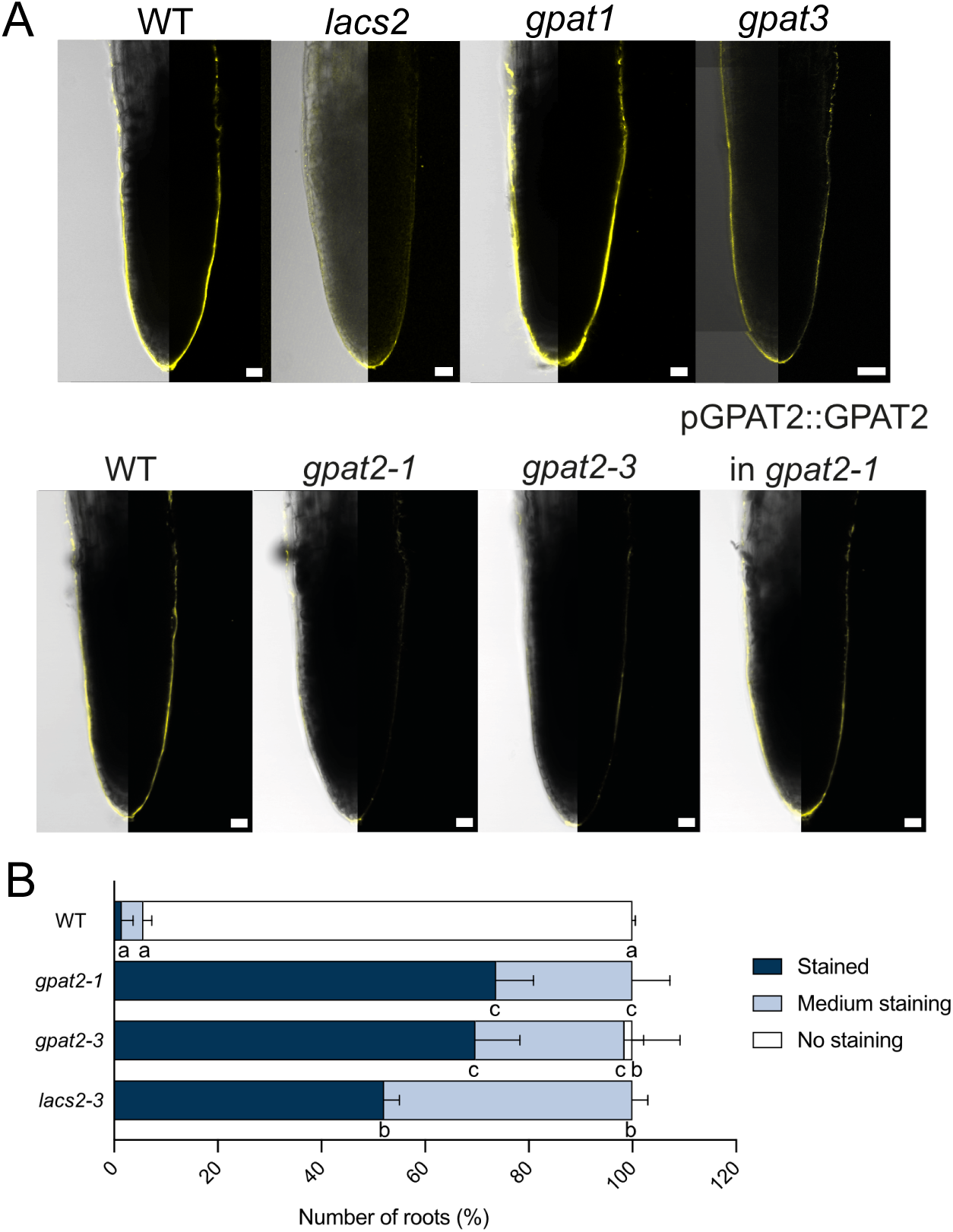
Characterization of cuticle at the root cap of the primary root in different genotypes. (A) Polyester deposition in the RCC of primary roots of different genotypes as visualized by Fluorol yellow staining. Median views of the root cap of 2-day-old seedlings are shown (on the left, overlay bright field and fluorescence; on the right, fluorescence only). Scale bars = 20 µm. (B) Assessment of the permeability of the root cap cuticle of the primary root by Toluidine Blue staining in different *gpat2* mutant alleles in comparison to the *lacs2* mutant and the wild type (WT). 20 to 30 roots were tested per replicates. Values represent the means of two repetitions and significant differences to wild type were assessed by ANOVA. Letters represent statistically significant differences between groups (p < 0.05).

## STAR METHODS

### KEY RESCOURCES TABLE

KEY RESOURCES TABLE
| REAGENT or RESOURCE | SOURCE | IDENTIFIER |
| --- | --- | --- |
| Antibodies |  |  |
| Monoclonal antibody to xylogalacturonan | Plantprobes | LM8 |
| Anti-Rat IgG–FITC antibody produced in goat | Sigma-Aldrich | Cat#F6257 |
| Chemicals, Peptides, and Recombinant Proteins |  |  |
| Fluorol Yellow 088 | Sigma-Aldrich | Cat#F5520 |
| Aniline Blue diammonium salt | Sigma-Aldrich | Cat#415049 |
| Toluidine Blue | Sigma-Aldrich | Cat#89640 |
| Fluorescein diacetate | Sigma-Aldrich | Cat#F7378 |
| ω-pentadecalacton | Sigma-Aldrich | Cat#W2840009 |
| Methylheptadecanoate | Sigma-Aldrich | Cat#H4515 |
| N, O- bis(trimethylsilyl)-trifluoroacetamide with trimethylchlorosilane (BSTFA) | Sigma-Aldrich | Cat#15238 |
| Pyridine | Sigma-Aldrich | Cat#270970 |
| Methyl acetate | Sigma-Aldrich | Cat#296996 |
| Sodium methoxide | Sigma-Aldrich | Cat#156256 |
| Dichloromethane | Sigma-Aldrich | Cat#650463 |
| Calcofluor white (Fluorescent brightener 28) | Polysciences | Cat#4359 |
| Xylitol | Sigma-Aldrich | Cat#X3375 |
| Sodium deoxycholate | Sigma-Aldrich | Cat#30970 |
| Urea | Sigma-Aldrich | Cat#51456 |
| Potassium hexacyanoferrate(II) trihydrate | Sigma-Aldrich | Cat#60280 |
| Low viscosity Embedding media Spurr | EMS | Cat#14300 |
| Osmium tetroxide 4% | EMS | Cat#19150 |
| Experimental Models: Organisms/Strains |  |  |
| *Arabidopsis: gpat2-1* | Yang et al., 2012 | SALK-118230 |
| *Arabidopsis: gpat2-3* | This study | SALK-64530 |
| *Arabidopsis: gpat3-2* | This study | SALK-139115 |
| *Arabidopsis: lacs2-3* | Bessire et al., 2007 | N/A |
| *Arabidopsis: gpat1* | Yang et al. 2012, Jia et al. 2024 | SALK-052352 |
| *Arabidopsis: pGPAT1::NLS-GFP-GUS* | This study | Transgenic Col-0 |
| *Arabidopsis: pGPAT2::NLS-GFP-GUS* | This study | Transgenic Col-0 |
| *Arabidopsis: pGPAT3::NLS-GFP-GUS* | This study | Transgenic Col-0 |
| *Arabidopsis: pGPAT5::NLS-GFP-GUS* | This study | Transgenic Col-0 |
| *Arabidopsis: pGPAT6::NLS-GFP-GUS* | This study | Transgenic Col-0 |
| *Arabidopsis: pGPAT7::NLS-GFP-GUS* | This study | Transgenic Col-0 |
| *Arabidopsis: pGPAT2::GPAT2* | This study | *gpat2-1* |
| Oligonucleotides |  |  |
| For all oligonucleotides used for genotyping and cloning | See Table S2 | N/A |
| Recombinant DNA |  |  |
| All recombinant DNA needed for the generation of transgenic lines are described in the subsection of the Star Methods - Method details- Generation of constructs | This paper | N/A |
| Software and Algorithms |  |  |
| Fiji | Schindelin et al., 2012 | https://fiji.sc/# |
| Rstudio | R Development Core Team, 2005 | https://www.rstudio.com/ |

### CONTACT FOR REAGENT AND RESOURCE SHARING

Further information and requests for resources and reagents should be directed to and will be fulfilled by the Lead contact Christiane Nawrath.

### EXPERIMENTAL MODEL AND SUBJECT DETAILS

#### Plant material

*Arabidopsis thaliana* accession Col-0 was used in this work. *Arabidopsis thaliana* mutants were already described: *lacs2-3* (Bessire et al., 2007), *gpat1* (SALK_052352; Yang et al. (2012), Jia et al. (2024)), *gpat2-1* (SALK_118230; Yang et al. (2012)); *gpat2-3* (SALK_64530; Jia et al. (2024); gpat3-2 (SALK_139115) were ordered at NASC, the European Arabidopsis Stock Center (Alonso et al., 2003). Gene numbers and genotyping primers are described in Table S1 and S2.

#### Growth conditions

Plants were grown under sterile conditions. Seeds were surface sterilized with chlorine gas. After 2-3 days of vernalization at 4°C, plants were grown on 1/2 MS (Murashige and Skoog, 500mg/l MES, pH 5.7), 0.7% agar at 22°C, under continuous light (100 µmol m^-2^ s^-1^). With the exception of the seedlings for polyester extraction, plants were grown vertically. For transformation and seed amplification plants were grown on soil under continuous light (100 µmol m^-2^ s^-1^) at 20°C and 65% humidity.

### METHOD DETAILS

#### Generation of constructs

To generate pENTRY L4-pGPAT1-R1, pENTRY L4-pGPAT2-R1, pENTRY L4-pGPAT3-R, pENTRY L4-pGPAT6-R1 and pENTRY L4-pGPAT7-R1, between 1.7 kb and 2.2 kb fragments upstream of each *GPAT* were amplified, respectively, and cloned into pDONR P4-P1 using KpnI and XbaI restriction site. All primers used are shown in Table S2. pENTRY L1-GPAT5-L2 was previously described (Naseer et al., 2012). To generate pENTRY-L1-GPAT2-L2, their respective gene was amplified and recombined into pDONR221. pGPAT2::NLS-GFP-GUS, pGPAT3::NLS-GFP-GUS, pGPAT5::NLS-GFP-GUS, pGPAT6::NLS-GFP-GUS, pGPAT7::NLS-GFP-GUS and pGPAT2::GPAT2 were generated by recombining the corresponding entry clones into the pFR7m24GW (Red seeds selection) using the Gateway Technology (Lifesciences). All constructs were transformed in *Agrobacterium tumefaciens* and then in Col-0 using the floral dip method (Clough and Bent, 1998) at the exception of pGPAT2::GPAT2 which was transformed in *gpat2-1*.

#### Cell wall polyester staining and Permeability assays

For visualization of cell wall polyesters, 10-20 roots were stained with Fluorol Yellow 088 (Berhin et al., 2019). Permeability assay using toluidine blue were done following the established protocol from Berhin et al., 2019 with slight modifications: 20-30 roots were staining for 30 sec with Toluidine Blue (0.05%/0.01% Tween 20), rinsed twice with water and evaluated immediately under the Axio Zoom V16 microscope (Zeiss) coupled to an Axiocam 512 Color camera. The experiment was repeated once.

For studying the permeability of the cuticle at the lateral root, with fluorescein diacetate (FDA) (5ug/ml in 0.5 x MS) (Barberon et al., 2016), FDA was applied on 25 lateral roots of the same developmental stage originating from 3 independent experiments following Berhin et al. (2019) protocol.

#### Root growth assays

Lateral root emergence was induced on 5-day-old seedlings by turning the plate of 90°C (Voss et al., 2015) and stages of lateral root emergence were evaluated after 42 h, using mutant roots having a comparable length to WT (Berhin et al, 2019) using a Leica DM5000B microscope. Experiments were repeated independently three times (n=20-30) and a representative data set is presented.

To examine the shape of the lateral root primordia, seedlings were fixed, cleared, and stained with Calcofluor following the method described by Ursache et al. (2018).

#### Immunofluorescence labeling

The protocol of Durand et al. (2009) was followed to label mucilage, using the LM8 antibody to detect xylogalacturonan-associated epitopes. The binding was visualized with a fluorescein isothiocyanate (FITC)-conjugated goat anti-rat antibody. A minimum of 30 roots of each genotype was studied.

#### Fluorescence Microscopy

Most fluorescent microscopy studies were performed on the confocal laser-scanning microscope ZEISS 700 with an excitation at 488 nm and detection with BP 490-555 nm for GFP, FY, FDA and LM8, and respectively at 555 nm and LP 640 nm for propidium iodide (PI). PI (10µg/mL) was used for staining the cell wall of the roots during the study of gene expression by direct application on the slide. Calcofluor staining was studied on the confocal ZEISS LSM 880 Airyscan with an excitation at 405 nm and detection at 425-475 nm. Red seeds selection was performed under the stereomicroscope Leica 6000 equipped with a DSR filter.

#### Transmission electron microscopy

The protocol and the microscope for transmission electron microscopy was identical than in Berhin et al. (2019). The ultrastructure of the root cap cell wall and cuticle was investigated over the entire length of the root cap in 2-3 independent root tip preparations per genotype and representative pictures were taken.

#### Chemical analyses

The full protocol for the determination of ester-bond lipids previously described in Berhin et al, (2019). 3 independent experiments were performed with 3-4 replicates for each genotype, respectively, and a representative data set is presented. The amounts of unsubstituted C16 and C18 fatty acids were not evaluated because of their omnipresence in the plant and in the environment.

### QUANTIFICATION AND STATISTICAL ANALYSIS

For the chemical analyses of the cutin composition, presented values are the mean ± standard deviation. Students T-test analyses were performed to highlight differences between WT and other genotypes. Asterisks illustrate the p-value: p < 0.001 is ***, p < 0.01 is ** and p < 0.05 is *.

Number of repetitions and replicates are mentioned for each experiment in the method details.

---

**Table S1.** Genes used in this study.

| Gene number | Gene name |
| --- | --- |
| AT1G01610 | *GPAT1* |
| AT1G02390 | *GPAT2* |
| AT4G01950 | *GPAT3* |
| AT4G00400 | *GPAT4* |
| AT3G11430 | *GPAT5* |
| AT2G38110 | *GPAT6* |
| AT5G06090 | *GPAT7* |
| AT4G00400 | *GPAT8* |
| AT1G49430 | *LACS2* |

**Table S2.**
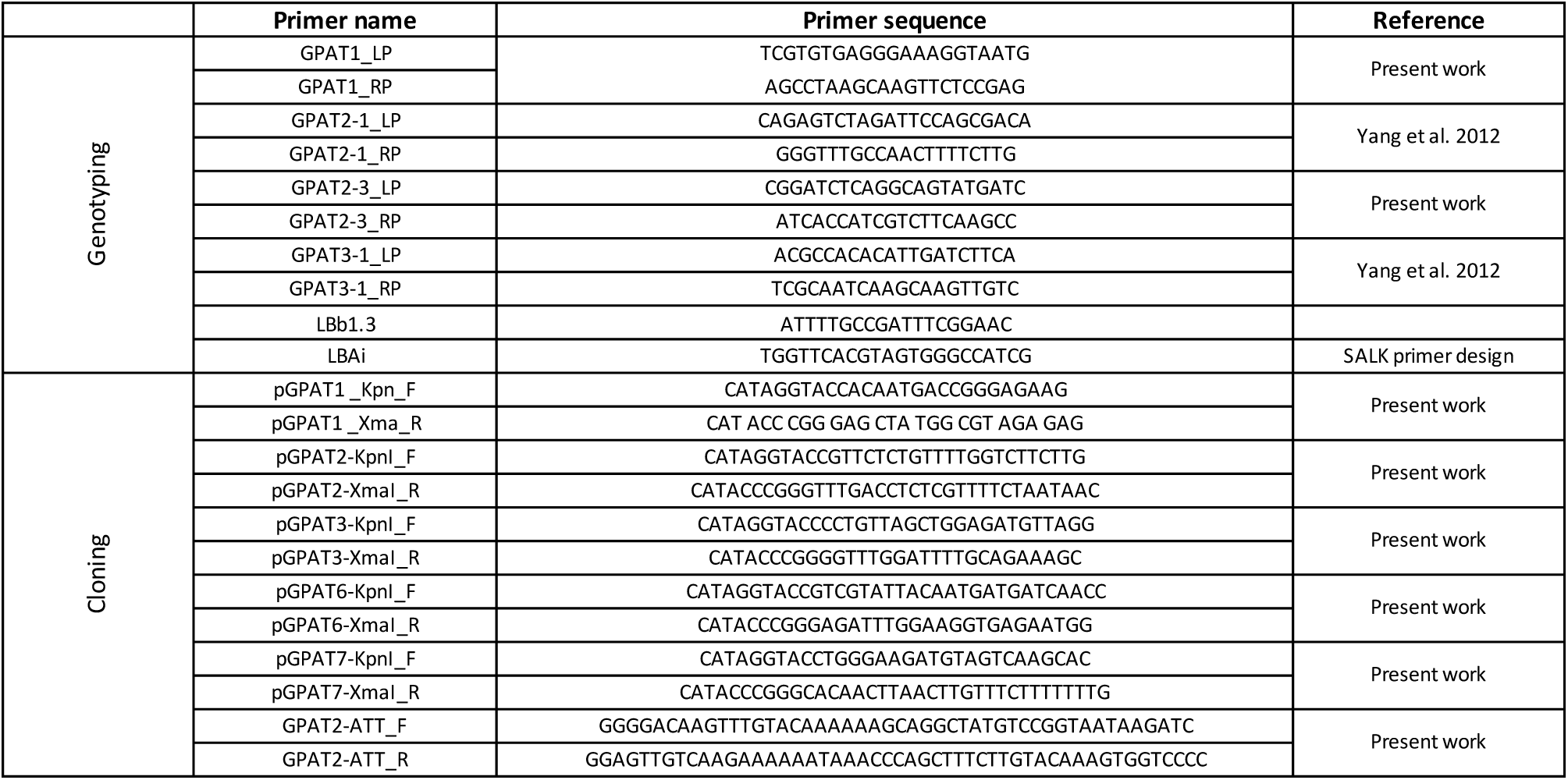
Primers used in this study.

## Notes

### Competing Interest Statement

The authors have declared no competing interest.

